# Clustering of Circular Consensus Sequences: Accurate Error Correction and Assembly of Single Molecule Real-Time Reads from Multiplexed Amplicon Libraries

**DOI:** 10.1101/236893

**Authors:** Felix Francis, Michael D. Dumas, Scott B. Davis, Randall J. Wisser

## Abstract

**BACKGROUND:** Targeted resequencing with high-throughput sequencing (HTS) platforms can be used to efficiently interrogate the genomes of large numbers of individuals. A critical challenge for research and applications using HTS data, especially from long-read platforms, is errors arising from technological limits and bioinformatic algorithms.

**RESULTS:** A single molecule real-time (SMRT) sequencing-error correction and assembly pipeline, C3S-LAA, was developed for libraries of pooled amplicons. By uniquely leveraging the structure of SMRT sequence data (comprised of multiple low quality subreads from which higher quality circular consensus sequences are formed) to cluster raw reads, C3S-LAA produced accurate consensus sequences and assemblies of overlapping amplicons from single sample and multiplexed libraries. In contrast, despite read depths in excess of 100X per amplicon, the standard long amplicon analysis module from Pacific Biosciences generated unexpected numbers of amplicon sequences with substantial inaccuracies in the consensus sequences. A bootstrap analysis showed that the C3S-LAA pipeline per se was effective at removing bioinformatic sources of error, but in rare cases a read depth of nearly 400X was not sufficient to overcome minor but systematic errors inherent to amplification or sequencing.

**CONCLUSIONS:** C3S-LAA uses a novel processing algorithm for SMRT amplicon-sequence data that produces accurate consensus sequences and local sequence assemblies. The community standard long amplicon analysis module from Pacific Biosciences is prone to substantial errors that raise concerns about findings based on this pipeline. The method developed here removed this confounding bioinformatics source of error, allowing for the identification of limited instances of errors due to DNA amplification or sequencing.

## Background

High-throughput sequencing (HTS) platforms have revolutionized the study of genomes and genomic variation. However, HTS platforms and analysis of HTS data generate errors in base calling (1). Even perfectly accurate sequence reads may be improperly assembled or incorrectly aligned to a reference sequence when read lengths are too short. The consequence of such errors can lead to incorrect results and misleading conclusions in a variety of settings ranging from scientific investigation (2) to clinical diagnostics (3).

Single molecule real-time (SMRT) sequencing by Pacific Biosciences (PacBio) generates long-read data, which, once error corrected [raw SMRT sequence data has a median error rate of ≈ 15% (4)], can help to produce complete de *novo* assemblies and accurate alignments to a reference genome. SMRT sequencing also exhibits relatively little sequence coverage bias, allowing regions of the genome with large differences in sequence complexity to be fully traversed (4). Therefore, SMRT sequencing facilitates assembly, resequencing, haplotype phasing, characterization of isoforms and structural variation, etc., all of which are more prone to errors with “short-read” data (5).

For resequencing applications, SMRT sequencing of tiled amplicons allows kilobase or larger-scale target regions of a genome to be sequenced at great depth, providing the opportunity to generate highly accurate, consensus assemblies (6). In combination with molecular barcoding, sequencing of multiplexed amplicon libraries facilitates studies across broad biological disciplines (7–11). However, such studies can be affected by confounding sources of errors arising during library preparation, sequencing and data analysis (12). Isolating the sources and types of errors is crucial to progress in the development of sequencing technologies, sequence analysis methods and interpretation of sequence data.

Several computational pipelines have been developed for automated processing and analysis of amplicon sequence data produced on different HTS platforms, such as PyroNoise (13), mothur (13) and Long Amplicon Analysis (LAA) (14). LAA is the standard pipeline for analysis of SMRT sequence data from amplicon libraries. LAA uses a “coarse clustering” approach to group raw reads according to pairwise similarity estimated from BLASR alignments (15). The Quiver consensus calling framework (16) is then used to generate an error-corrected consensus sequence for each cluster. When we first used LAA to process amplicon sequences as part of a previous study (6), several of the consensus sequences outputted by LAA were incorrect. We found that clustering of high quality circular consensus sequences (i.e. clustering of CCS reads, which we refer to as C3S) to group the corresponding raw read data prior to performing analysis with Quiver recovered all of the expected sequences with high fidelity. Here, we investigated this further and present a new, open-source pipeline for processing tiled amplicon resequence data from multiplexed libraries.

## Methods

### Sequence data

PacBio sequence data from two amplicon libraries, a single sample library (SRX2880716) and a multiplex sample library (SRX3474979), were used for this study. SMRTbell libraries were constructed according to PacBio’s amplicon library protocol (17). Sequencing was performed on a Pacbio RS II instrument with one SMRT Cell used for each library, using P6/C4 chemistry with a 6-hour movie. SMRTbell library preparation and sequencing was carried out by the University of Delaware Sequencing and Genotyping Center (Newark, DE).

Sequence data from the single sample library was from a previous study (6) and was comprised of nine amplicons, which were amplified from the maize inbred line B73. The maximum expected amplicon size was 4,954 bp, such that the raw reads, which had a mean length of 23,794 bp, consisted of an average of approximately nine subreads per amplicon (Table S1). The multiplex sample library produced for this study was comprised of a tiling path of six amplicons spanning æ 23,000 bp of the maize genome, which were amplified from six different maize inbred lines (B73, CML277, Hp301, Mo17, P39, Tx303). The primer pairs used for the multiplex library had distinct symmetric barcodes for each sample and amplicon, along with a shared 5’ GTTAG padding sequence (Table S2). The maximum expected amplicon size was 7,752 bp and the raw reads consisted of an average of nine subreads per amplicon (Table S1).

### Clustering of circular consensus sequences for long amplicon analysis

A cluster and assembly pipeline was developed in which raw reads are clustered based on circular consensus sequences (CCS) prior to running error correction with Quiver. We refer to this pipeline as C3S-LAA, for Clustering of Circular Consensus Sequence (C3S) Long Amplicon Analysis.

Clustering is performed as follows. The reads of insert protocol in SMRT Portal is used to generate CCS reads (run settings: minimum of 1 subread at 90% CCS read accuracy). These are higher quality sequences formed from the corresponding raw reads based on their multiple subreads. Therefore, the CCS reads are used to cluster the data. Clustering is performed by a simple match function that identifies CCS reads containing both the forward and reverse primer sequences for each amplicon (considering the sense and antisense primer sequences). From this, a list of CCS read identifiers belonging to each amplicon cluster is produced. This list is then used to subset the corresponding raw reads, using the whitelist option in LAA, such that Quiver-based consensus calling (16) occurs on only the raw reads belonging to a given amplicon-specific cluster. Consensus sequences formed from clusters comprised of fewer than 100 subreads were eliminated when all available reads were used; this setting was adjusted to 0 for evaluation of accuracy (see below). The pipeline can be used to perform one-level clustering for non-barcoded amplicon libraries or two-level clustering for barcoded amplicon libraries. Because barcodes or other sequences may precede the primer sequence and may vary in length, the primer search space was designed as a user input parameter, which, for this study, was set to 21 bases at both the ends of the sequence.

The pipeline proceeds to an assembly step (Figure 1). The C3S-LAA consensus sequences are automatically merged into a Multi-FASTA format file and assembled (per barcode if barcoding is used) using Minimus based on the overlap-layout-consensus paradigm (18). To trim extraneous sequences (e.g. padding or barcodes) for downstream analysis, a user input parameter (trim_bp) is specified to remove the corresponding number of bases from each end of the consensus sequences while writing them to the FASTA file. The assembly is then carried out among all trimmed consensus sequences, and mismatches between any two overlapping sequences are represented as Ns in the assembly sequence. Where there are more than two overlapping sequences with mismatches, the most frequent base will be represented in the assembly. In the case of barcoded sequencing libraries, the assembly is carried out separately for each barcode.

**Figure 1.**
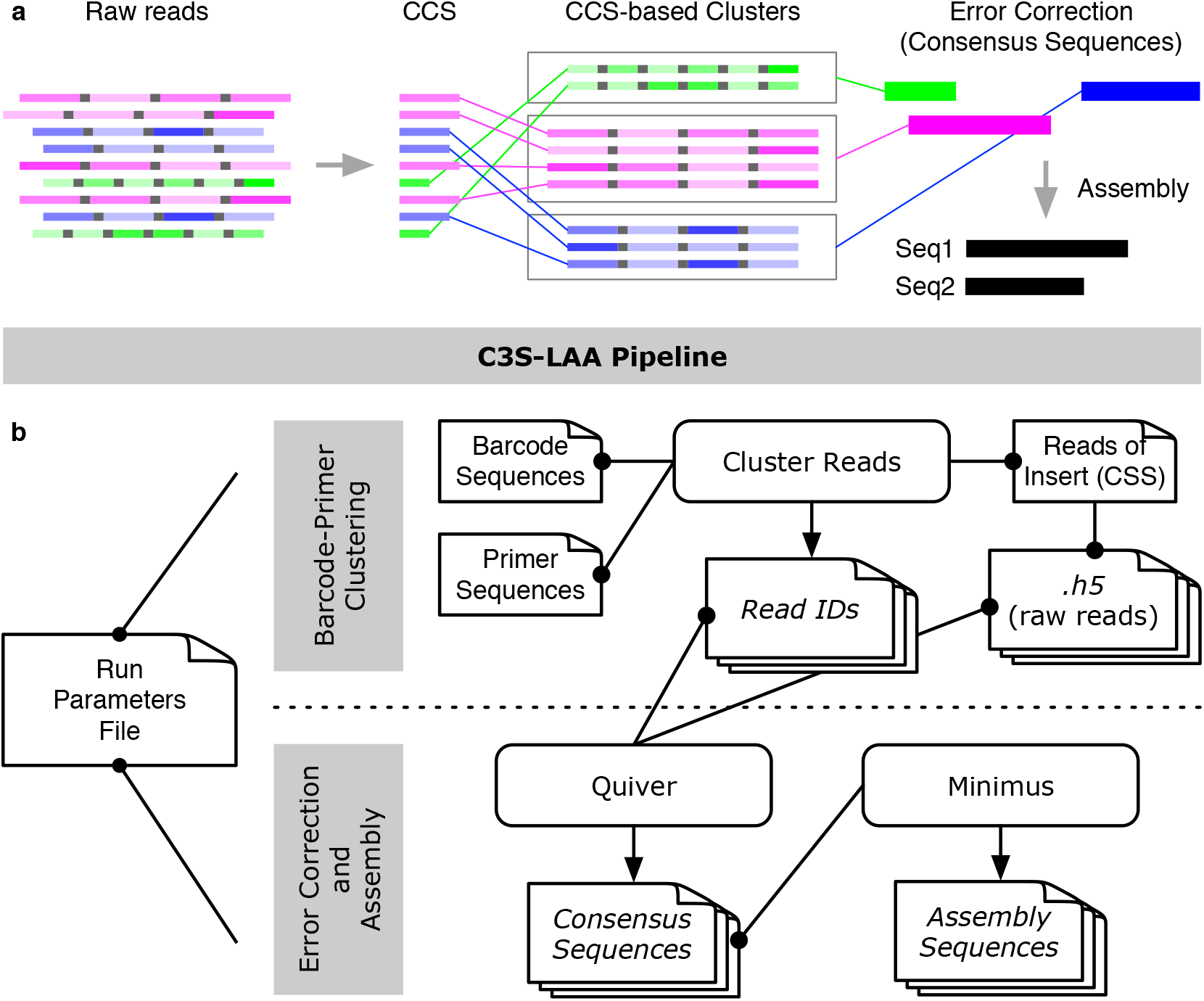
Graphical representation of the C3S-LAA process and pipeline. (a) Raw reads comprised of multiple subreads are depicted for three different amplicons [green, fuchsia and blue boxes; different shades of color are used to portray variable subread sequence qualities (darker shading portrays higher quality)]. Subreads are separated by a shared adapter sequence (grey boxes). The higher quality CCS read for each raw read is used to cluster the corresponding raw reads into CCS-based cluster groups. Error correction is performed per CCS-based cluster, producing top quality consequences sequences, followed by assembly of any overlapping consensus sequences. (b) A single run parameters file is used by all components of the pipeline. The grey highlighted rectangles represent two main steps of C3S-LAA. (i) Using the CCS reads generated by the SMRT analysis reads of insert protocol, C3S clusters the raw reads according to each barcode-primer pair combination, producing files of read identifiers to whitelist the corresponding raw reads. (ii) Raw read clusters are passed to Quiver to generate amplicon-specific consensus sequences, which are then passed to Minimus for sequence assembly. Rectangles with folded corners represent single files or multiple files (depicted as stacks of files) and those with rounded edges represent scripts and tools. Arrows indicates output files that are generated. Connecting lines with dots at one end depict input files, with the dot corresponding to the source data for the connected script or tool.

### Evaluating the accuracy of C3S-LAA

First, evaluation of the performance of LAA was carried out on the sequence data from the single sample library. LAA v1 was run on SMRT Portal, using the following settings: minimum subread length: 2000 bp; maximum number of subreads: 2000 (default); ignore primer sequence when clustering: 0 bp (default); trim ends of sequences: 0 bp (default); provide only the most supported sequences: 0 (0=disabled filter; default); coarse cluster subreads by gene family: yes (default); phase alleles: no; split results from each barcode into independent output files: no; barcode: no. The minimum subread length was reduced from the default value of 3000 bp to 2000 bp since the sequencing library had one amplicon of 3330 bp, such that any partial sequences may also be considered. Phasing of alleles was not used since the amplicons were produced from homozygous individuals (inbred lines). The resulting LAA consensus sequences were aligned using BLASTn (19) to the B73 v3 reference genome of maize (20). YASS (21) was used to align the incorrect (partial matches) consensus sequences formed by LAA to their expected amplicon sequence using the following score parameter settings: Scoring matrix (match: +5, transversion: -4, transition: -3, composition bias correction: -4), Gap costs (opening: -16, extension: -4), E-value threshold: 10 and X-drop threshold: 30.

The same sequence data from above was also processed using C3S-LAA. In addition, the relationship between subread depth and the accuracy of consensus sequence construction as well as assembly was evaluated for the output from C3S-LAA. For each amplicon, sample sets of 1,2,3,…40 CCS read identifiers were randomly selected with replacement from among the eight amplicons. Using the corresponding raw reads of each CCS read set, C3S-LAA was used to create consensus sequences per amplicon cluster and assemblies from the corresponding group of consensus sequences belonging to a sampled CCS read set. This was repeated 25 times, such that a total of 8000 consensus sequences were generated in addition to the corresponding Minimus assemblies. BLASTn alignments with the B73 v3 reference genome were used to determine the map location and compute the percent identity for each of the amplicon-specific consensus sequences and corresponding assembly sequences. From these alignments, the number of mismatches and gaps were also recorded to characterize the types of errors present in the sequences. For each cluster of sequences, the number of subreads used to derive the consensus sequence was recorded. The minimum number of subread counts for a set of overlapping amplicons that produced an assembly was used as the number of subreads for that assembly.

The performance of LAA versus C3S-LAA was also evaluated using the multiplex sample library. LAA was used to generate consensus sequences under the same settings indicated above, with an additional selection of the barcode demultiplexing option. Since the amplicons were barcoded using PacBio’s standard barcodes, the default pre-set in SMRT Portal pointing to PacBio barcodes with padding in the reference directory was used. C3S-LAA was used to perform two-level clustering of the CCS reads, using the primer and barcode sequence information. A search space of 121 bp was used for identifying barcode-primer sequences in order to cluster the CCS reads. Since one of the lines (B73) has a reference genome available, LAA and C3S-LAA consensus sequences associated with B73 were aligned using BLASTn to the B73 v3 reference genome. The C3S-LAA assembly for B73 was also compared to this reference.

## Results and Discussion

PacBio SMRT sequence data from a pooled library of long-range PCR amplicons was previously produced and used for this study (6). The data was processed with PacBio’s LAA protocol under default settings using all of the raw read data. This did not produce a consensus sequence for all of the expected amplicons and included several artifactual sequences (Table 1). Dot-plot visualization of the alignments of these seven incorrect consensus sequences with the reference sequence indicated the presence of spurious inverted duplications in six out of the seven consensus sequences and a truncated alignment for the one remaining sequence (Figure S1).

**Table 1.** Comparison of LAA and C3S-LAA consensus sequences for B73.

| Library type<sup>1</sup> | Method | Number of consensus sequences | Complete match<sup>2</sup> (100% identity) | Truncated match (100% identity) | Partial match (<100% identity) |
| --- | --- | --- | --- | --- | --- |
| Single | LAA | 14 | 7 | 1 | 6 |
| Single | C3S-LAA | 9 | 9 | 0 | 0 |
| Multiplex | LAA | 8 | 4 | 1 | 3 |
| Multiplex | C3S-LAA | 6 | 5 | 0 | 1 |

<sup>1</sup>The single library had nine expected consensus sequences, whereas the multiplex library had six expected consensus sequences.
<sup>2</sup>For the multiplex sample library, the B73 v3 assembly contained a gap relative to one of the five amplicon sequences. This gap was filled in the latest B73 v4 release.

The above errors led us to inspecting LAA, which uses a custom algorithm based on the raw read data to pre-cluster similar sequences for analysis by Quiver. However, PacBio raw reads are notoriously poor in quality and overlapping or repetitive sequences could be present, either of which may cause errors in cluster formation (our speculation based on results presented below). Moreover, the primer sequences used to produce the amplicons in a library are not considered. Therefore, we hypothesized that using the higher quality CCS reads to group the corresponding raw reads into amplicon-specific clusters based on the expected primer sequences would improve the consensus sequence analysis. A bioinformatic pipeline, C3S-LAA, was developed to carry out such clustering (Figure 1). The divide and conquer principle of C3S-LAA simplifies the determination of consensus sequences by only operating on raw reads for which there is a high degree of certainty that they were derived from the same origin.

Indeed, our results indicated that C3S-LAA rectified the errors generated by the standard LAA protocol. For a typical use case, where all the reads from a sequencing library are used, C3S-LAA could resolve and accurately call the consensus sequence for every amplicon in the single sample library with no extraneous sequences (Table 1). In contrast, more than half of the consensus sequences generated by LAA had truncated or partial matches to the reference genome, and LAA could only fully resolve six of the nine amplicons in the single sample library. Based on these results, we recommend C3S-LAA for analysis of PacBio amplicon sequence data. Moreover, the C3S concept may be used in other situations where some portion of the sequences are known in advance.

For tiled amplicon resequencing, C3S-LAA can also be used to assemble any overlapping segments that may exist among the consensus sequences outputted for a given genotype (Figure 1). We bootstrapped the read data from the single sample library to examine the accuracy of the assemblies, as well as the underlying consensus sequences, produced by C3S-LAA as a function of subread depth. All C3S-LAA alignments of the resulting amplicon consensus and assembly sequences mapped to the expected target region. The minimum subread depth from which amplicon-clustered consensus sequences were outputted by LAA was 21, which corresponds to « 2 CCS reads for our 3-5 kb amplicons (mean number of passes was 9.39; Table S1). Accuracy of the consensus sequences from bootstrapped samples of amplicon-clustered data was generally high, with accuracies ranging from of 99.72-100% (Figure 2a). By extension, Minimus assemblies of these consensus sequences were similarly accurate (Figure 2b). Despite an increase in accuracy with subread depth, not all of the bootstrap replications from high CCS sample depths included completely accurate consensus sequences or assemblies, and even at a subread depth of nearly 400X some bootstrap samples included imperfect assemblies (Figure 3). This was primarily due to a specific error in one locus (locus_6_7045710_7052049) that was observed among some of the bootstrapped samples at different CCS sampling depths (rare instances of locus_1_25390617_25396540 also showed minor inaccuracies). For instance, at a CCS sample depth of 40, the consensus sequence for locus_6_7045710_7052049 contained a 2 bp insertion in two of the 25 bootstrap samples. This same type of insertion error occurred for both loci and was embedded within homopolymeric regions of the sequences (Figure 4), indicating this was due to PCR or sequencing and not the pipeline per se. Among all the assemblies generated from bootstrapping (n = 3787), errors in the form of insertions, deletions and single nucleotides contributed to 66.7, 17.2 and 16.1% of the total errors respectively.

**Figure 2.**
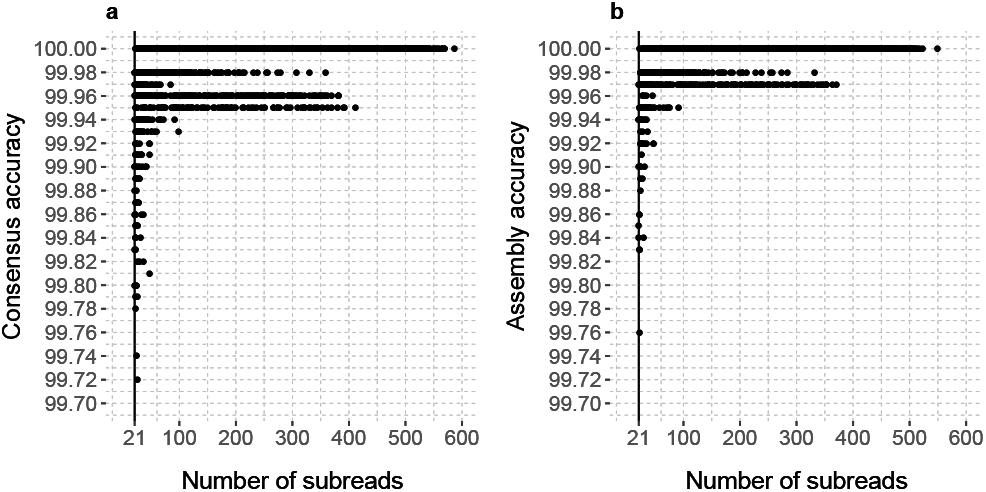
Sequence accuracy as a function of subread depth. (a) Accuracy of consensus and (b) assembly sequences. Data from all the amplicons were pooled together to evaluate the consensus calling accuracy as a function of depth of coverage of SMRT raw reads. The vertical line shows the minimum read depth of the consensus sequences used for assemblies.

**Figure 3.**
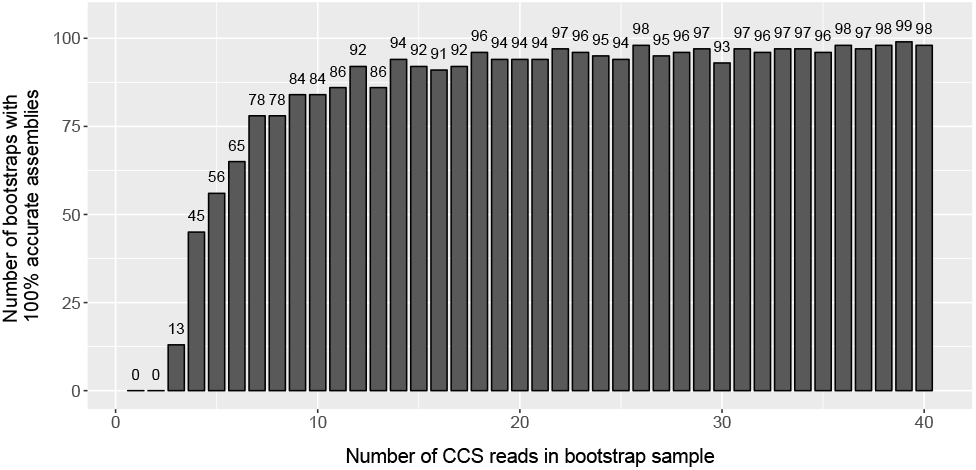
Total number of accurate bootstrap assemblies per CCS sample size. At each level of the CCS read depth sample (1–40), the figure shows the total number of bootstrapped assemblies that were 100% identical to the reference sequence. This was determined forthe fourtarget regions (25 bootstrap assemblies at each of 4 loci, giving rise to a maximum of 100 on the x-axis) formed from the consensus sequences among the eight overlapping amplicons.

**Figure 4.**
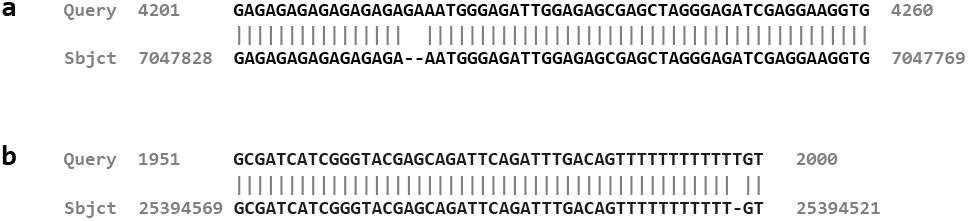
Sequence alignment highlighting a recurring insertion error in some bootstrap samples. The alignment corresponds to the consensus sequence for a part of the amplicon from (a) locus_6_7045710_7052049 (Query) and (b) locus_1_25390617_25396540 (Query) on maize chromosome 6 and 1 respectively compared to the B73 v3 reference sequence (Sbjct).

For the multiplex sample library, the number of consensus sequences formed by LAA differed from the expected number for four of the six samples, and LAA generated consensus sequences for barcodes that were not used to make the library (Table 2). In contrast, C3S-LAA produced the exact number of expected consensus sequences per sample and per barcode. As with the single sample library, comparing the B73-barcode derived consensus sequences to the B73 reference genome showed substantial errors in the consensus sequences from LAA but not C3S-LAA, where LAA only resolved four of the amplicons from B73 (Table 1); the one C3S-LAA consensus sequence with an imperfect match was due to two separate 1 bp insertions embedded within homopolymeric regions. Another C3S-LAA consensus sequence aligned to the expected region of chromosome 1 with 100% identity but spanned a 531 bp assembly gap in the reference genome. This gap was filled in the latest release of the B73 reference genome [v4; (22)] which matched perfectly to the C3S-LAA consensus sequence. None of the other results were changed when using the B73 v4 reference sequence. C3S-LAA also produced assemblies for each sample from the corresponding set of consensus sequences. The 23,300 bp C3S-LAA assembly for B73 differed from the expected B73 reference genome sequence only by the differences indicated above.

**Table 2.** The number of consensus sequences generated from the multiplex library, following barcode demultiplexing.

| Sample<sup>1</sup> | Barcode ID<sup>2</sup> | LAA | C3S-LAA |
| --- | --- | --- | --- |
| B73 | 32 | 8 | 6 |
| CML277 | 35 | 6 | 6 |
| Hp301 | 31 | 6 | 6 |
| Mo17 | 20 | 7 | 6 |
| P39 | 2 | 7 | 6 |
| Tx303 | 4 | 7 | 6 |
| N/A | 8 | 7 | 0 |
| N/A | 23 | 5 | 0 |
| N/A | 49 | 1 | 0 |
| N/A | 82 | 2 | 0 |
| N/A | 85 | 1 | 0 |
| N/A | 91 | 6 | 0 |
| N/A | 92 | 3 | 0 |

<sup>1</sup>No samples were associated with the N/A barcode.
<sup>2</sup>Identifier for PacBio barcodes.

C3S-LAA clearly outperformed LAA for the data examined in this study. We have observed the same performance using C3S-LAA on data from a multiplex library of another 21 individuals amplified across multiple overlapping amplicons (not shown). Nevertheless, a potential limitation of C3S-LAA is that it requires the CCS reads have both the barcode and primer sequences intact. Accuracy of CCS reads is a function of the number of subreads (23). Thus, for very long amplicons where one or a few subreads are sequenced, reliance on CSS reads will limit the number of sequences used from the available data. It may be possible to use a less stringent clustering algorithm, however, the fragment lengths of most amplicon libraries are expected to be well below the current and increasingly long read lengths of PacBio data, such that fairly accurate CCS reads would be available for clustering. C3S-LAA is expected to be applicable for SMRT sequence data of amplicon libraries or where flanking sequences can be predefined. C3S-LAA was developed as part of an extension to tiled amplicon resequencing projects facilitated by ThermoAlign (6) and is released under an open source license.

## Conclusions

This study shows that CCS-facilitated clustering of raw reads vastly improves the analysis of SMRT sequence data. This method directs error correction and consensus sequence analysis to be performed only on sequences derived from the same amplicon and sample, leading to accurate consensus sequences and local sequence assemblies. The community standard LAA module could not resolve all of the expected amplicons in a library and generated several spurious consensus sequences during barcode demulitplexing and sequence clustering. LAA v1 uses BLASR (15) for pairwise alignment of all reads, which are then clustered based on their similarity using a Markov Model (R. Lleras, Pacific Biosciences of California, Inc., pers. comm.). Given that the the underlying principle of LAA and C3S-LAA are essentially the same — use clustering to group reads from which consensus sequences should be formed — but only C3S-LAA produces correct output, indicates that the clustering algorithm of LAA is prone to error. This release of C3S-LAA provides users with a more accurate processing pipeline for SMRT sequence data, which addresses a critical gap in the analysis of amplicon sequence data.

## Availability

The code for C3S-LAA is released under an MIT open source license at: https://github.com/drmaize/C3S-LAA.

## Author’s contributions

RJW guided the study. FF and RJW conceived the design principles for C3S-LAA. MDD and SBD produced the sequence data. FF developed the code and executed the computational analysis and program optimization, which iterated based on feedback from RJW. FF and RJW wrote the manuscript.

## Acknowledgements

This work was supported by the U.S. NSF Plant Genome Research Program IOS-1127076. We thank Dr. Karol Miaskiewicz at the Delaware Biotechnology Institute for assistance with using BIOMIX, a high performance computing cluster. BIOMIX was supported by Delaware INBRE grant NIH/NIGMS GM103446 and a number of investigators who have contributed nodes to the cluster.

